# A novel computational approach for genome-wide prediction of small RNAs in bacteria

**DOI:** 10.1101/011668

**Authors:** LI Lei, Hoi Shan Kwan

**Author notes:** Present address: Institute for Molecular Infection Biology, University of Würzburg, Germany (LI Lei). Email Addresses: LL HSK.

## Abstract

Small regulatory RNAs (sRNAs) are the most abundant post-transcriptional regulators in bacteria. They serve ubiquitous roles that control nearly every aspects of bacterial physiology. Identification of important features from sRNAs sequences will guide the computational prediction of new sRNA sequences for a better understanding of the pervasive sRNA-mediated regulation in bacteria. In this study, we have performed systematic analyses of many sequence and structural features that are possibly related to sRNA properties and identified a subset of significant features that effectively discriminate sRNAs sequences from random sequences. we then used a neural network model that integrated these subfeatures on unlabeled testing datasets, and it had achieved a 92.2% recall and 89.8% specificity. Finally, we applied this prediction model for genome-wide identification of sRNAs-encoded genes using a sliding-window approach. We recovered multiple known sRNAs and hundreds of predicted new sRNAs. These candidate novel sRNAs deserve extensive study to better understand the sRNA-mediated regulatory network in bacteria.

## Introduction

Bacteria small regulatory RNAs (sRNAs) are a class of independently functional transcripts. Most sRNAs do not translate into proteins (Storz et al., 2011b). They are heterogeneous in size with around 50 and 300bp long. sRNAs are generally classified as antisense RNAs, which are encoded on the opposite strand of coding sequences and trans-encoded sRNA, which are only partially complementary to their targets, with frequently mediated by RNA-binding protein Hfq. There is also a kind of sRNAs, which are interacting with proteins by mimicking other nucleic acids. In any case, the interactions of these highly structured sRNAs and their targets could result in activation or inhibition of the target genes function at either post-transcriptional or translational levels or both.

Although only a few sRNAs have been functionally well characterized, it is widely believed that sRNAs are involved in nearly every aspect of bacterial physiology. sRNAs such as Qrr1-5 in *Vibrio cholera* are involved in biofilm formation and motility, and control the bacterial transitioning from the planktonic to surface-associated lifestyle (Chambers and Sauer, 2013). Several sRNAs are often identified in pathogenic islands and regulate virulence in some pathogens. SprD, a sRNA identified in *Staphylococcus aureus,* negatively regulates the expression of immunoglobulin-binding protein (Sbi) and further impairs the host immune responses (Chabelskaya et al., 2010). They are often synthesized under specific conditions and act as important regulators of gene expression in response to external environmental stimuli such as low iron medium, oxidative stress, and elevated glucose-phosphate levels (Thomason et al., 2012).

Traditionally, bacterial sRNAs can be identified by size fractionation of total RNAs on denaturing gels and excision of specific bands for further studies (Majdalani et al., 1998) . However, the use of direct sequencing for identification of RNA molecules encountered many obstacles, for example, the amount of RNA should be abundant enough for visualization and the experimental screening is time-consuming. The emergence of bacterial genome sequences provides possibilities for global identification of sRNAs using computational searches. Because sRNAs mostly have independent transcriptional units and are a class of structural RNAs, current identification methods were generally based on a) primary sequence or secondary structural conservation, or b) peculiar genomic features such as Rho-independent terminators, promoter sequence and predicted secondary structures. These features have been effectively utilized and integrated in various programs (Argaman et al., 2012; Storz et al., 2011b). RNAz integrates the features based on structural conservation and thermodynamic stability to identify structural non-coding RNAs (Altuvia et al., 1997; Thomason and Storz, 2010). QRNA identify structural RNAs by searching of covariance patterns in multiple sequence alignments (Babitzke and Romeo, 2007; Papenfort et al., 2013). sRNAPredict is the first program that specifically identifies sRNAs based on a combination of transcription signals and sequence conservation (Argaman et al., 2012; Lu et al., 2011).

Recently, whole genome expression profiling methods such as differential transcriptome sequencing (Altuvia et al., 1997; Sharma et al., 2010) and tiling arrays has advanced the global discovery and quantification of sRNAs in bacteria (Sittka et al., 2008; Thomason et al., 2012). Many of these methods detect expression signals from non-protein-coding regions, which led to the identification of numerous candidate sRNAs. Co-immunoprecipitation (coIP) sequencing of specific RNA-binding proteins enabled further functional characterization of sRNAs interactions. Sittka et al (2008) compared the RNA transcriptomes from the RNA coIP with epitope-tagged Hfq protein and control coIP, and found more than 30 novel Hfq-associated sRNAs (Lu et al., 2011; Majdalani et al., 1998).

Although there are many existing tools at our disposal, a recent study indicated that the current computational prediction methods for novel sRNAs yielded low precision (6%-12%) and sensitivity (20%-49%) (Argaman et al., 2012; Livny et al., 2006). Algorithms such as sRNAPredict (Altuvia et al., 1997; Livny et al., 2008), SIPHT (Papenfort et al., 2013; Sridhar et al., 2010), and sRNAscanner (Gruber et al., 2010; Lu et al., 2011) are not effective for predicting sRNAs without provision of distinct sequence features such as promoter sequences or rho-independent terminators. This is a drawback for detecting sRNAs especially for those with non-intrinsic terminators. There are several other algorithms based on structure RNA conservation properties such as RNAz (Elena Rivas, 2001; Sharma et al., 2010), eQRNA (Lu et al., 2011; Sittka et al., 2008). These algorithms rely on the compensatory changes of multiple sequence alignment, however, the evolutionary distances that lie outside an optimal range could cause poor performance (Lu et al., 2011; Ott et al., 2012). Several other methods are solely based on sequence conservation such as NAPP (Chao et al., 2012; Livny et al., 2006), and are not helpful for the identification of ‘orphan’ sRNAs. More importantly, nearly all sRNAs finders only search within intergenic region, excluding sRNAs in the UTR region, which has recently been found as an sRNA rich depository (Livny et al., 2008; Tjaden, 2008). In addition, sRNAs are likely to be expressed under rare conditions or are hard to distinguish from UTR transcripts such as 5’ UTR. These issues underscore the systematic detection of sRNAs in bacterial genomes still remains an fruitful and challenging problem, and a highly reliable novel approach is needed.

In this study, we have analyzed many sequences and structural features possibly related to sRNA properties. In total, 27 features including 10 sequence and 17 structural features have been identified that show significant discrimination between sRNAs with randomly shuffled sequences. Furthermore a neural network with Adaboost machine learning approach was applied to unlabeled test datasets. This approach has received over 92.2% recall and 89.8% specificity to accurately classify sRNA sequences from negative sequences. To our knowledge, this is the first systematic analysis of sequence and structural features within sRNAs sequences, which would serve as a valuable guide for novel sRNA discovery. We have further developed a neural network model by extracting multiple valuable features among known sRNA sequences for the prediction of sRNAs. This approach had a high performance in classifying sRNA sequences from random sequences in the unlabeled datasets. We have used this model for genome-wide identification of sRNAs-encoded sequences using a sliding-window approach and found that several known sRNAs that were not included in the training datasets can be successfully recovered as well as hundreds of predicted new sRNAs. These studies suggest there could be many sRNAs unidentified even in the typical genomes.

## Results

### 1. Description-based sequence features cannot resolve the existence of sRNAs

Previous work demonstrated that current computational methods for predicting sRNAs have low performance, however, evaluating the contribution of each feature has not been considered. In this study, we compiled the largest collection of experimentally validated sRNA sequences from the BSRD database, and redundant sequences were removed using MCL (Enright et al., 2002) resulting in 699 unique clusters (see Material and Methods). Training sRNAs were then randomly selected from each cluster. These sRNAs were experimentally identified by various methods including Northern blot, RT-PCR and direct sequencing (Li et al., 2013). We examined the length distribution of these sRNAs sequences and it was found that 96.48% of them have sequence size smaller than 500bp, which is consistent with the definition of sRNA (Storz et al., 2011b).

Since the transcriptional signals were often used to predict sRNAs, we first determined whether most sRNAs sequences possessed these features. We analyzed all the sRNA upstream sequence to detect the signals of promoter sequences including -35 region, -10 region and transcriptional start site using a combination of PPP and NNPP by setting the threshold with a range of 0.8 to 1. At the same time, we searched for the putative terminators within the downstream of sRNA sequences using TranstermHP. We detected the promoter sequence signals in 31% of the upstream of sRNAs, and rho-independent terminators were present in 61.1% of the downstream of sRNAs. Only a minor percentage (12.3%) contain both promoter and terminator sequences, whereas nearly half of sRNAs did not have either promoter or terminator motifs (Table 1).

**Table 1.** Promoter and terminator consensus sequences found

| Consensus<sup>a</sup> | N | % |
| --- | --- | --- |
| promoter POS; terminator POS | 86 | 12.3 |
| promoter POS; terminator NEG | 131 | 18.7 |
| promoter NEG, terminator POS | 141 | 20.2 |
| promoter NEG, terminator NEG | 341 | 48.8 |
| Total | 699 | 100 |

<sup>a</sup>Presence (POS); Absence (NEG).

To further determine the relationship between various types of sRNAs and transcriptional signals of sRNAs, we have further analyzed the Rho-independent terminators in two groups of trans-encoded sRNAs and cis-encoded antisense RNAs using unpaired t-test, we found cis-encoded antisense RNA had significantly low (unpaired t test: P < 0.0001) rho-independent terminators compared to trans-encoded antisense sRNA. This result suggests that most of cis-encoded antisense RNAs may require non-intrinsic terminators such as rho-dependent terminators (Peters et al., 2009).

I have further analyzed the sRNA sequence conservation patterns using RNAz (Gruber et al., 2010), which is a program for predicting non-coding RNAs with consideration of the secondary conservation and thermodynamic stability. Among 956 validated sRNAs, 604 sRNAs were predicted to have the structural features, and 352 sRNAs (36.8%) were not detected. 145 sRNAs were not predicted due to the missing of homologs within the NCBI non-redundant database. The others proved to be false positive.

Taken together, the above analysis suggests that the current computational approaches for sRNA prediction that rely on transcription signals or conservation profile are unlikely to effectively characterize the sRNAs sequences. While combining the traditional features with cross-species conservation may improve the performance to some extent, it is still challenged by not discovering orphan sRNAs. Recent data also showed that if the evolutionary distance lies outside of the optimal range, this could cause bias to the detection of sRNAs (Lu et al., 2011).

### 2. Analysis of sRNAs primary sequence reveals new features

I have investigated a few primary sequence features, including conservation, non-coding properties and sequence composition frequency to determine whether sRNAs show distinct sequence features from negative datasets. First, conservation is a widely used feature to identify sRNAs-encoded genes, and evolutionary conservation often suggests functional importance. However the sequence conservation from sRNAs has not been systematically explored before. Here we have conducted a comparative genomics-based search for sRNA homolog sequences in the NCBI non-redundant nucleotide sequences to determine the extent of sequence conservation in other organisms. Figure S1 shows that 52.4% of sRNAs are conserved within less than eight genomes, which suggest that sRNAs are not broadly conserved, but only conserved in closely related genomes.

Second, sRNA transcripts previously thought to be purely non-coding may in fact encode small proteins (Vanderpool et al., 2011). To assess the protein-coding potential of the sRNA data, we utilized the program CPC (Kong et al., 2007) to measure the protein-coding potential of sRNAs. We compared these results with randomly selected protein-coding transcripts from Uniprot database. Figure S2 shows that sRNAs have a very different coding potential from protein-coding genes (Wilcoxon signed rank test: P < 3.368e-171). Thus, at least at the sequence level, the sRNAs sequences do not appear to have similar protein-coding sequences.

Third, we expect that the evolutionary forces that shape the diversity of sRNAs could lead to a unique sequence distribution than other elements. To support this hypothesis, the mono-and di-nucleotide frequency on sRNAs and randomly selected protein sequences from Uniprot database were analyzed (UniProt Consortium, 2013). For the mono-nucleotide frequency, we found that the nucleotide frequency of G and U are most significant, as showed by P-value lower than 0.0001. Likewise, %AC, %CA, %CC, %CG, %CU, %UA, %UG,%UU, %GA, %GC and %GU is the significant dinucleotide with %UU ranked the most significant one (unpaired t test: P = 1.63e-55) (Figure 2).

**Figure 2.**
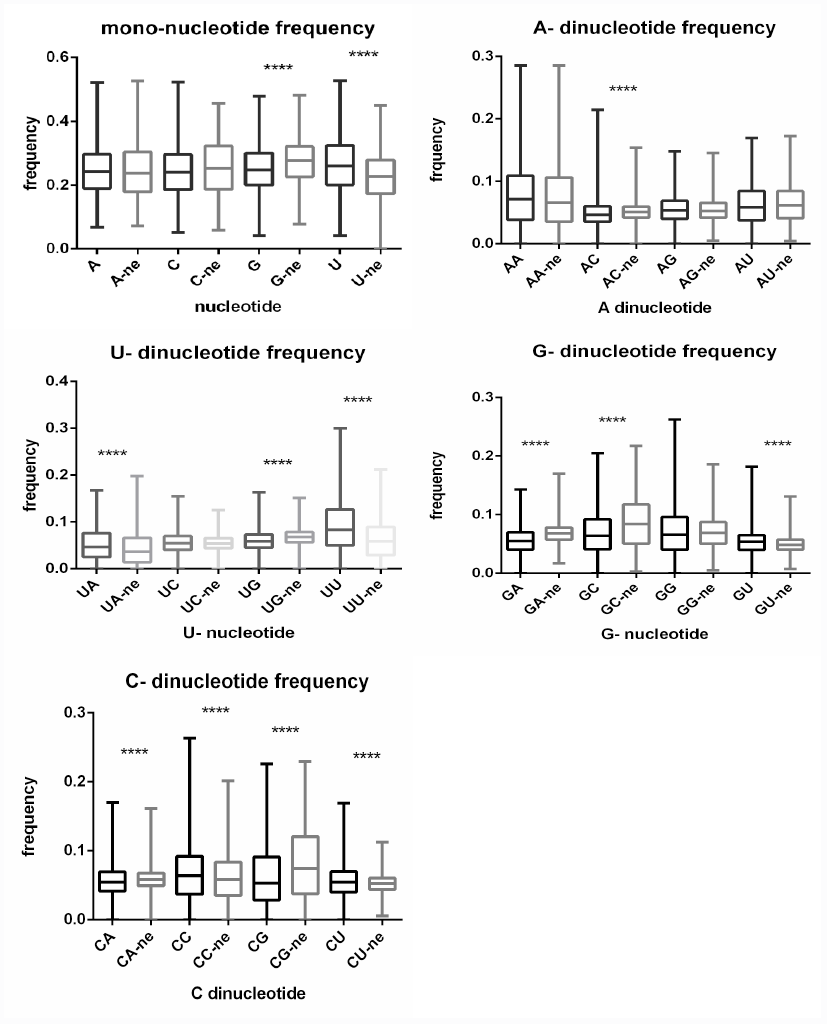
The mono-and di-nucleotides frequency of sRNAs. (The dinucleotide frequency from positive and negative datasets are compared and **** indicate the P < 0.0001 using unpaired t test. The boxplot in black color indicate the calculation from positive datasets, the boxplot in grey color indicate the calculation from negative datasets)

### 3. Evaluation of individual secondary structures features

In addition to examine primary sequence features, we also investigated a large set of secondary structure features (Table 2), which may potentially be beneficial for the characterization of the properties of the sRNA-encoded genes. These features were chosen mainly based on previous studies focusing on structural RNA prediction and many other features that possibly helped. These include the minimum free energy folding statistics including minimum free energy, the P value for representing the percent of the number of random di-shuffled sequence with MFE larger than the original sequence and the Z-score describing the standard deviation between the di-shuffled sequence and the original sequence.

**Table 2.** Features used in this study.

| Features | Numbers | Description |
| --- | --- | --- |
| **sRNA primary sequence** |  |  |
| Single nucleotide sequence | 4 | Frequency of each nucleotide (A, T, C, G) |
| Dinucleotide frequency | 16 | Frequency of each dinucleotide |
| **MFE Folding statistics** |  |  |
| MFE | 1 | Normalised minimum free energy (dG) |
| P-value | 1 | The significant of the permutation test |
| Z-score | 1 | Number of standard deviation that a sequence deviates from shuffled sequences |
| **Ensemble statistics** |  |  |
| Free energy | 2 | Average free energy of the ensemble and the frequency of MFE structure in the ensemble |
| Structure Clustering features | 14 | Statistics of structure clustering in the Boltzmann ensemble |
| Ensemble diversity | 1 | Base pair distance between the MFE and suboptimal structures in the ensemble |
| Shannon entropy | 1 | Reliability of the secondary structure prediction |
| **Structural statistics** |  |  |
| Stem | 3 | Total stem number, each stem number and the average stem length |
| Loop | 13 | Average number and total number of loop, internal loop, multiple loop |
| Bulges | 4 | Total bulges number, each bulge number and the average stem length |

Thermodynamics stability was considered to be the contravention features of structural RNAs. It was first proposed by Maizel that structural RNAs are more thermodynamically stable than random sequences (Le et al., 1988). However, further evidence suggests that the secondary structure alone is not statistically significant to detect structural RNAs. (Kavanaugh and Dietrich, 2009). In spite of this, these features were found to be able to discriminate microRNA precursors from other genetic elements (Bonnet et al., 2004). So to explore whether sRNAs could have distinct thermodynamic stability compared to random di-shuffled sequences, we selected experimentally validated sRNA sequences and their randomly shuffled sequences and then calculated the minimum free energy of sRNAs, P-value, which represents as the percent of numbers of randomly shuffled sequences in 1000 permutation tests, and Z-score, which is described as the standard deviation from the normal randomly shuffled sequences. It was found that although the number of P-value larger than 0.05 is nearly the same as the number of P-value smaller than 0.05, the distribution of P-value and Z-score is generally lower than that of random sequences (unpaired t test: P < 0.0001) (Table S1). This suggests that these two folding statistics based on minimum free energy are significant for sRNA discovery. Despite the fact that P-value and Z-score are somewhat correlated (Freyhult et al., 2005), integrating these two values have the advantages over just using one of them. However, folding free energies are not good signals for detection of sRNAs (unpaired t test: P = 0.371) because the expected free energies will generally decrease linearly with the size of the sequence.

We have identified features based on a minimum free energy based structure, and subsequently analyzed a series of features based on an ensemble of sRNA structures, where ensemble refers to a collection of non-crossing suboptimal structures of the same RNA sequences (Chan and Ding, 2008). We included numerous ensemble and clustering-based features for evaluating global folding reliability and clustering quality among all optimal and suboptimal structures such as the ensemble diversity, the compactness of each cluster and the between- and within- cluster sum of squares.

We evaluated whether these features can be distinguishable with negative datasets. All the ensembles-based features are normalized by the sequence length for statistics evaluation. From S1, most of ensemble features have highly significant distinction from random sequences, which suggest the robustness of these features. The overall compactness, which is a measure of the global density among all the structures, are generally lower than random sequences, while the number of clusters are also in general lower, which suggests that sRNA secondary structure tends to form fewer clusters and be more densely clustered than random sequences, which is in agreement with Tran et al (2009)’s previous conclusion. Interestingly, in these densely clustered sequences, it also shows a large number of high-frequency base-pairs. Taken together, the examined ensemble and clustering features have revealed that sRNAs possesses distinct global folding and clustering quality compared with random sequences.

In addition, to further determine whether sRNAs shows any different structural features compared to random sequences, we used 18 structural statistics, which were previously described by Tran et al. (2009). In table S2, most of the structural component features were different from random sequences excepting the average number of loop. We found that sRNAs in general have compact stem secondary structure (Table S2), which shows fewer stem number with larger average nucleotide numbers in each stem structure. Further investigating the loop properties of sRNA sequences found that sRNAs tended to have larger loops than random sequences, however, the average nucleotide of sRNAs in a single loop region is not significant.

### 4. Training and evaluation of sRNAs classifiers

As the number of non-sRNA sequences were much larger than that of positive datasets, 1398 sets of sequences were randomly selected from the shuffled negative pool to construct the negative test datasets. These negative datasets were combined with training sRNA sequences to form the final testing datasets. To determine the contribution of each feature, we have used a feature ranking method to identify these sub features from the full 41 features. We found that while many features show distinction between positive and negative datasets, the power of these features varies in different levels. In total, 17 sub features were identified with the average merit larger than 0.06 using Info Gain evaluator implementation with 10-fold cross validation in WEKA (Frank et al., 2004) (Table 3). We further examined these sub features and found that most of these features have overlapped with Tran’s list (Tran et al., 2009). They identified ten features with the mean of AUROC larger than 0.6, and these ten features are all with average merit larger than 0.06 in my result, although the ranking order is different. However, three new MFE folding statistics features used in this study ranked as the best performance among all these features with average merits of 0.396 and 0.258 separately. We have not identified any structural statistical features except for the average of the stem in each stem component.

**Table 3.** The contribution of each feature evaluated with Information Gain ranker

|  |  |  |  |
| --- | --- | --- | --- |
| Features | Average merit | Average rank |  |
| p-value | $0.396 \pm 0.027$ | 1 | $\pm 0$ |
| z-score | $0.258 \pm 0.005$ | 2 | $\pm 0$ |
| maxcompactness | $0.152 \pm 0.008$ | 3 | $\pm 0$ |
| overallcompactness | $0.137 \pm 0.007$ | 4 | $\pm 0$ |
| ave_bpdist_mfe_ensemble | $0.122 \pm 0.004$ | 5 | $\pm 0$ |
| avecompactness | $0.115 \pm 0.007$ | 6.4 | $\pm 0.66$ |
| stem_ave | $0.111 \pm 0.004$ | 7.1 | $\pm 0.94$ |
| compactnesslargest | $0.111 \pm 0.006$ | 7.6 | $\pm 0.49$ |
| bss_point | $0.099 \pm 0.004$ | 9.1 | $\pm 0.3$ |
| bss | $0.097 \pm 0.006$ | 9.8 | $\pm 0.6$ |
| ensemble_diversity | $0.083 \pm 0.004$ | 11.4 | $\pm 0.49$ |
| nclusters | $0.082 \pm 0.005$ | 11.6 | $\pm 0.49$ |
| num_hifreq_ensemble | $0.069 \pm 0.002$ | 13.6 | $\pm 0.92$ |
| wss | $0.067 \pm 0.003$ | 13.9 | $\pm 0.3$ |
| wss_point | $0.067 \pm 0.004$ | 14.5 | $\pm 0.81$ |
| dG | $0.062 \pm 0.003$ | 16.2 | $\pm 0.4$ |

In this study, we applied three different machine learning approaches to determine whether they can efficiently classify positive and negative sequences from the pooled datasets. These three machine learning classifiers included a random forest classifier, a decision tree classifier and a neural network classifier. We have made a meta approach with each classifier combining with Adaboost, which was evidenced to improve machine learning performance (Leclercq et al., 2013). The datasets were evaluated using 10-fold crossvalidation section. Each classifier was trained on the full set of 41 features or on the selected of 17 features based on feature ranking with a score larger than 0.06. From table 3.6, the neural network with AdaBoost using selected features received the best performance, with a sensitivity of 92.2%, and a specificity of 89.8%. Interestingly, the neural network with AdaBoost on the full set of features, however, produces slightly weaker results. Random forest with Adaboost generates slightly lower sensitivity, but with poor specificity. The C4.5 decision tree with J48 model received the worst performance (Table 4).

**Table 4.**
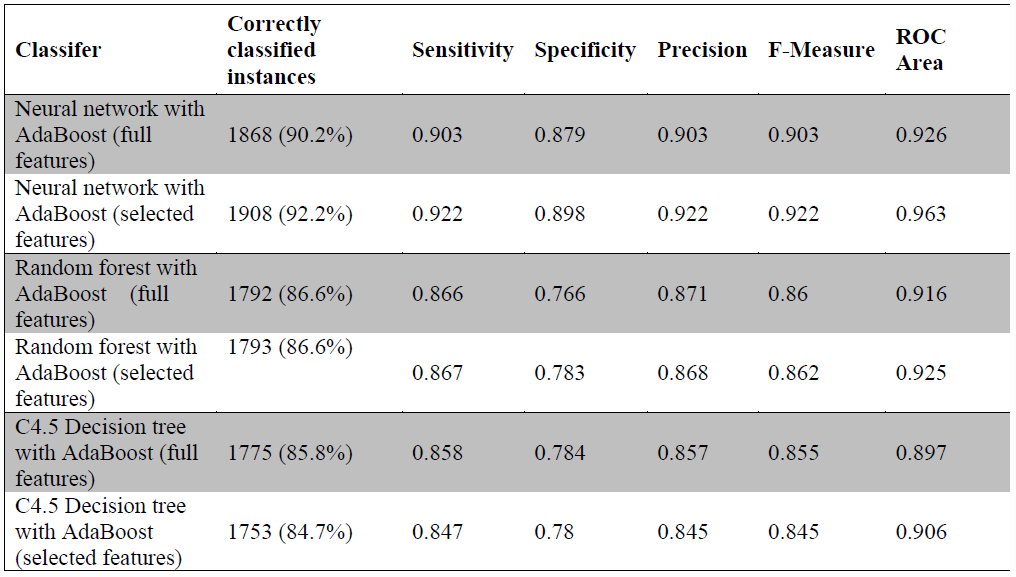
Results of various classifers using 10-fold cross validation.

### 5. The workflow of sRNADeep*

Our goal was to detect the sRNA-encoded genes in a genome-wide fashion. For this purpose, we collected four genome sequences including two gram-positive and two gram-negative strains. We used a fixed window size of 160bp with 50 step size. The sliding-windows were scanned in the forward and reverse strand of the genome sequences. The resulting sequence in each window was compared to non-redundant protein sequences in NCBI using BLASTx. The windows with no homolog or an alignment length lower than 80bp to known protein sequences were kept for further analysis. For the case of *Escherichia coli,* which have a genome size of 4,639,675 bp. Sliding-window approaches have generated the number of 92,791 windows in the forward and in reverse strand. These small windows were further filtered, leaving 11,329 unannotated windows in the forward and 11,299 unannotated in the reverse strand (Table 5).

**Table 5.** The partitioned genomes using sliding-windows approach

|  | Gram-negative |  | Gram-positive |  |
| --- | --- | --- | --- | --- |
| Strains | *Escherichia coli*<br>MG1655 | *Salmonella*<br>Typhimurium<br>LT2 | *Listeria monocytogenes*<br>EGD-e | *Streptococcus agalactiae*<br>NEM316 |
| Genome size | 4639675 bp | 4857432 bp | 2944528 bp | 2211485 bp |
| Windows (forward) | 92791 | 97146 | 58888 | 44227 |
| Windows (reverse) | 92791 | 97146 | 58888 | 44227 |
| Unannotated windows (forward) | 11329 | 13005 | 7157 | 5829 |
| Unannotated windows (reverse) | 11299 | 13043 | 7122 | 5849 |

Thirty seven features previously described including 17 structural features and 20 sequence features were calculated and integrated. We made a two-step neural network model training to avoid the biased models built by two unpaired negative datasets. The first step was to train the prediction model on structural features in each window size. The second step was to train the sequence model training from sRNA sequence statistics in each window size again. Only the positive results from both two steps were considered as the candidate sRNAs (Figure 3).

**Figure 3.**
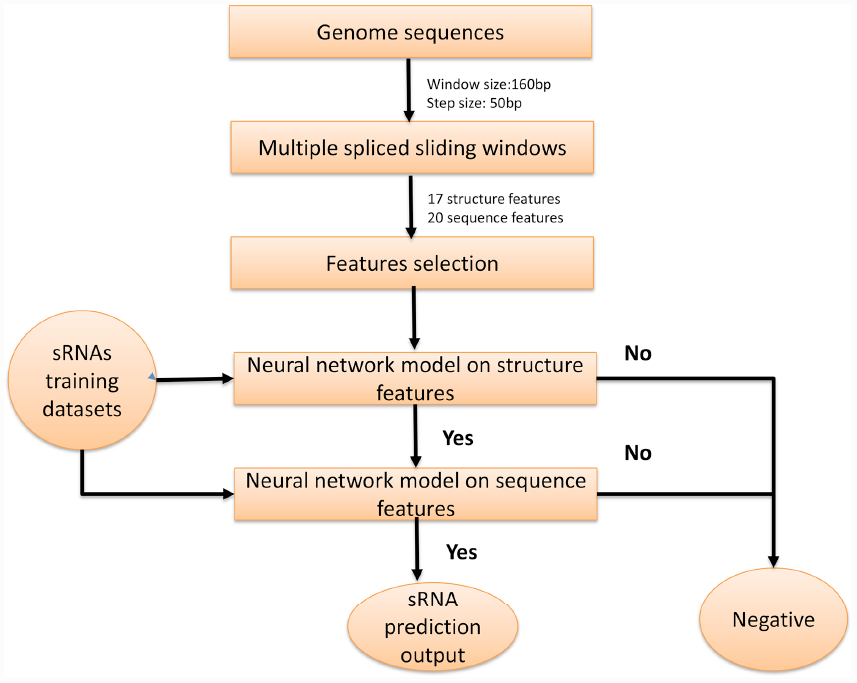
The workflow for sRNADeep*.

### 6. Genome-wide prediction of sRNAs using sRNADeep*

The candidate sRNAs location predicted with sRNADeep* were compared to known sRNAs sequences stored in the BSRD database to evaluate the number of known sRNAs recovered. To further identify the novel sRNAs from the candidate pool, the predicted windows were compared to open reading frames of each strain, and the windows overlap with these ORFs of their genomes were considered as the location of cis-encoded antisense RNAs with the remaining regarded as candidate trans-encoded sRNAs.

*Escherichia coli* is an intensively analyzed model organism, and 108 sRNAs were identified in the past decades. Of these known sRNAs, sRNADeep* could effectively identify 84 sRNAs, accounting for 77.8% of the total number (Table S3). A summary of the sensitivity (Sn) and positive prediction values (PPV) for the other methods is summaried in table 6. Compared with the previous methods, my approach received the lower candidate number, however it had with best performance. Results of sRNADeep* prediction using a series of selected structure and sequence-based features were promising given that we did not consider other transcriptional signals such as promoter and terminator elements, also the sequence covariance built by multiple sequence alignment frequently used by other programs. Although Tran et al. (2009) also used a de novo approach to predict the existence of sRNAs, they did not consider the thermodynamic features and used a different model training approach, which comes to nearly 37% lower sensitivity compared with my approach. We have also found 314 novel sRNAs, which had been missing in previous studies (Table S4). Further experiments such as northern blot or RT-PCR are needed to validate the existence of some sRNAs. Based on the nature of sRNAs, it is highly possible that there could be many more previously overlooked sRNAs in *Escherichia coli.*

**Table 6.** Comparison of prediction accuracies by different approaches for *Escherichia coli*

| Program | No. of prediction | Sn<sup>a</sup> | PPV<sup>b</sup> | References |
| --- | --- | --- | --- | --- |
| Carter | 563 | 0.3441 | 0.0568 | Carter et al., 2001 |
| Chen | 227 | 0.2903 | 0.1189 | Chen et al., 2002 |
| Rivas | 275 | 0.4086 | 0.1382 | Rivas et al., 2001 |
| Saestrom | 306 | 0.1183 | 0.0359 | Saetrom et al., 2005 |
| Wang | 420 | 0.0753 | 0.0167 | Wang et al., 2006 |
| Tran | 601 | 0.4086 | 0.0632 | Tran et al., 2009 |
| sRNADeep* | 398 | 0.778 | 0.211 | This study |

<sup>a</sup> the sensitivity of program
<sup>b</sup> the positive prediction value

*Salmonella* Typhimurium is another model gram-negative organism. There were 119 sRNAs recorded in BSRD. Of these sRNAs, sRNADeep* identified 87 sRNAs, accounting for 73.1% of known sRNAs (Table S5). Kroger et al (2012) used dRNA-Seq technology and identified multiple intergenic sRNAs. Interestingly, most of these sRNAs have also been detected by sRNADeep*. sRNADeep* not only identified intergenic sRNAs, but can also be can used to detect sRNAs in UTR region. DapZ is a Hfq-dependent sRNA that shared 3’ UTR and Rho-independent terminator regions with dapB CDS. Due to the fact that DapZ is in close proximity with mRNA sequences and possesses poorly conserved sequences with related organisms makes it previously escaped by all of computational methods (Chao et al., 2012). However, sRNADeep* has managed to detect the location of this sRNA.

sRNADeep* has been applied to gram-positive bacteria such as *Listeria monocytogenes* and *Streptococcus agelactiae. Listeria monocytogenes* is a common foodborne pathogen, which can result in brain and materno-fetal infections. Toledo-Arana et al (2009) had previously used tiling microarray to compare the transcriptomes of wild type and multiple mutants. They have identified more than 50 sRNAs, that may be associated with virulence. Oliver et al (2009) also used the RNA-Seq to compare the multiple transcriptomes between stationary phase cells and sigB mutant, and a total of 67 sRNAs were identified by using a combination of HMM model training (Oliver et al., 2009). Among these sRNAs identified from various sources of high throughput technology, sRNADeep* have recovered 93 sRNAs (76.2%) solely based on primary sequence. The application to *Streptococcus agalactiae* also yielded the average prediction sensitivity of 71.4%. The known sRNAs of *L. monocytogenes* and *S. agelactiae* identified by sRNADeep* are summarized in Table S6, Table S7, respectively.

## Discussion

In this project, we analyzed various sequences and structural features and have identified a subset of features that significantly discriminates sRNAs from their randomly shuffled sequences. We developed a neural network with Adaboost classifier for identification of sRNAs by integrating these informative features, which received high performance.

In this study, a sliding-window approach was also used to detect sRNA-encoded genes of the genomes of four bacteria. A window size of 160bp and step size of 50bp were used. Although it was arbitrary for fixing the sliding-windows, this approach has achieved the highest performance. The sliding-window approach was previously used for the genomewide screen of non-coding RNA. Tran et al. (2009) used three fixed windows size of 100, 120 and 160 nucleotide, which indicated the three peaks of their training datasets. However, in my training datasets, we did not identify other peaks except the 160bp. A possible reason is that the training sets they used were from various non-coding RNAs in bacteria including the riboswitches, and several other leader elements. Kavanaugh et al (2009) used an intelligent method, which investigated on multiple sliding windows with different step sizes based purely on single sequence folding energy, and have successfully identified several novel ncRNAs in *Saccharomyces cerevisiae.*

Although the prediction of small RNAs in bacteria has been extensively studied, and numerous programs have been developed. The computational prediction of sRNAs still resulted in in a low sensitivity and accuracy (Lu et al., 2011). The current predictions mostly rely on description-based method such as transcription factor binding site, promoter site or rho-independent terminators, or rely on comparative genomics approaches. Compared with previous approaches, the approach we developed does not rely on a prior knowledge of sequence covariation with closely related genomes, or the knowledge of transcriptional signals. It is simply based on a few sequence and structural features derived from primary sequence, and can be directly applied to any complete or partial bacterial genomes.

The discovery of sRNA sequences is of great importance. Firstly, although only a few sRNAs have been well characterized, it is now generally believed that sRNAs play a crucial regulatory role in bacterial physiology. Identifying novel sRNAs and further characterizing their binding targets will enhance our comprehensively understanding of the roles they played. Secondly, we are still unclear on why the abundant small molecules exist within diverse species. With the efficient prediction tools and comparative genomics analysis, we can give a glimpse into the evolution of sRNAs.

The novel approach complements a wide range of primary sequence features and secondary features that aim to identify sRNAs. Also the training datasets comes from BSRD, a repository for experimental validated sRNA sequences, which represent as the goldstandard dataset repository for sRNAs. This is also the first attempt to comprehensively analyze these experimental validated sRNAs sequences.

**Table 7.**
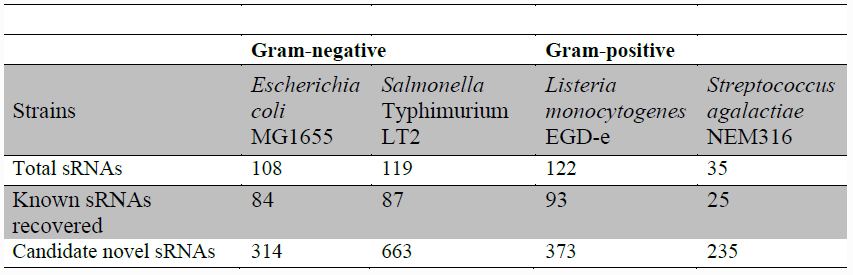
Summary result predicted by sRNADeep*

sRNADeep* doesn’ t need multiple sequence alignment or any algorithms for predicting candidate transcriptional signals, and is simply based on a few sequence and structural features derived from primary sequence and showed a more robust performance than previous computational approaches (Lu et al., 2011). The successful application of sRNADeep* to two gram-negative and two gram-positive bacteria suggests that it would be useful to annotate sRNAs regions in other gram-positive and gram-negative bacteria. sRNADeep* is a powerful tool for detection of small RNAs in the era of high throughput sequencing data. Computational prediction of sRNAs in genomic sequences would facilitate experimental studies such as expression profiling under various conditions, functional assay by deletion or mutagenesis, identification of interaction partners or structural analysis to extensively explore the sRNA-mediated regulatory network.

While this approach can be efficiently used for discriminating sRNAs compared to other random sequences, it cannot characterize the genomic boundaries of sRNAs. Future studies are needed to integrate transcription signals derived from known sRNAs or location information across the genome.

## Conclusion

In summary, we have made the following contributions and conclusions in this study (Figure 4).

**Figure 4.**
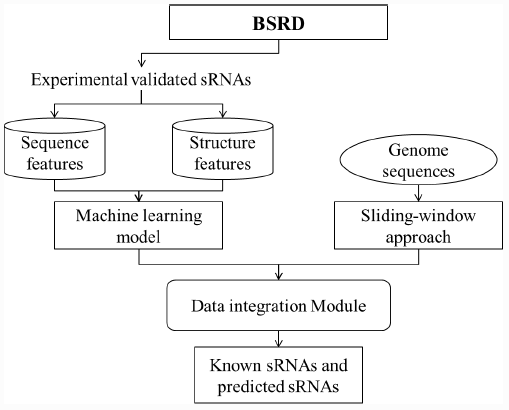
The summarized workflow of this study.

We have made systemic analysis of sequence and structural features with possible relations to sRNA properties and several novel features have been identified that show significant discrimination between sRNAs with random sequences. These features include three folding statistics (P-value, Z-score, and dG), and several ensemble-based features. We have further used a neural network based machine learning approach on unlabelled datasets, and this approach has received an overall result of 92.2% sensitivity and 89.8% specificity to accurately classify sRNA sequences from negative sequences.

Based on informative features identified above, we have further developed the updated version of sRNADeep, two-step neural network approach for annotating sRNAs from whole genome sequences using a sliding-window approach. An advantage of sRNADeep is that it only replies on the sequence itself, without considering the comparative genomics or other transcription signals. We have utilized the sRNADeep for a genome-wide identification of sRNAs. We found that over 70% known sRNAs were correctly predicted using sRNADeep, also hundreds of predicted new sRNAs were identified. This study may suggest that there could be many more unidentified sRNAs even in the typical genomes. We hope this study can provide new insights into the determinants of sRNAs sequences, and the BSRD database and the sRNADeep tools can provide useful resources to the research community of bacterial small regulatory RNAs.

## Material and Methods

### 1 Data source

All the experimentally validated sRNA sequences were retrieved from BSRD database (Cao et al., 2009; Sridhar et al., 2010). A total of 1095 sRNAs were collected and classified as trans-encoded sRNAs, cis-encoded sRNAs, and protein binding sRNAs. To remove redundant sequences within these datasets, these sequences were pairwisely compared to each other with an e-value set to 1e-2 using NCBI BLAST, then the Markov cluster (MCL) algorithm was applied to cluster similar sequences with the default inflation parameter (-I 2) (Busch et al., 2008; Gruber et al., 2010) . Finally 716 clusters were obtained and protein binding sRNAs, which have a distinct pattern, were manually removed. Then we randomly selected one sRNA from each grouped cluster and 699 sRNAs sequences were used for constructing positive control datasets. The sRNAs selected for analysis were summarized in Table S8.

Two negative datasets were also created. One negative training set of non-sRNAs was created by randomly selecting CDS sequences from Uniprot database (Elena Rivas, 2001; Tafer and Hofacker, 2008) using a python script. Each negative set is twice as large as its corresponding positive set. The second dataset is the generation of random shuffled 1000 permutations of sRNA sequences using ushuffle program (Lu et al., 2011; Storz et al., 2011a). These shuffled sequences preserved their mono-and di-nucleotides composition, due to the problems of stacked base-pairs (Ott et al., 2012; Sharma et al., 2011).

### 2 Primary sequence analysis

#### 2.1 Transcriptional signal analysis

Promoter sequences and transcription factor binding sites were predicted by two approaches. The first approach was to use the PPP web server, which determined the promoter sequences based on multiple trained HMM models (http://bioinformatics.biol.rug.nl/websoftware/ppp). The second method is using the Neural Network Promoter Prediction program (NNPP, www.fruitfly.org/seq_tools/promoter.html ). The principle of NNPP is to incorporate structural and compositional properties of the promoter sequence, and apply a machine learning-based approach to model these features. Intrinsic terminator sequences were identified using TranstermHP v2.09 (Chao et al., 2012; Tjaden et al., 2006).

#### 2.2 Coding potential analysis

The coding potential of sRNAs was analyzed using CPC program (Kong et al., 2007) with default parameters. Sequences of sRNAs were compared for similarities in Uniprot Ref90 database (meaning that the sequence similarity threshold of each cluster is larger than 90%). Then the pre-built model was applied to the sequence features extracted from sRNA sequences to predict the coding potential.

#### 2.3 Sequence-based nucleotide frequency

The four mono- (A, T, C, G) and 16 di-nucleotides (AA, AT, AC, AG, TA, TT, TC, TG, CA, CT, CC, CG, CA, CT, CC, CG, GA, GT, GC, GG) frequencies were calculated using a custom-made perl script.

#### 2.3 Secondary structure analysis

##### 2.3.1 RNA MFE folding statistics

(1) Normalized minimum free energy (dG). The normalized energy is defined as

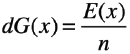

Where *E(x)* is the minimal free energy of a sequence (x), and n is the length of the sequence (x).

(2) P-value and Z-score

The Z-score compares the minimum free energy (MFE) of a sequence (x), to the distribution of MFE generated by randomly shuffling a sequence with the same mono-and di-nucletiode composition. The MFE of each sequence was calculated using the RNAfold program (Hofacker and Stadler, 2006). Each sequence was then shuffled 1000 times using the ushuffle program (Jiang et al., 2008). The mean and standard deviation was calculated for the resulting distribution. The Z-score was then calculated using the equation below. A P-value is also used to represent the significant of the permutation test.

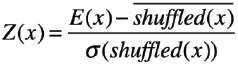

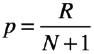

Where 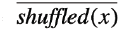 and denote the mean and standard deviation of the MFE of the shuffled sequence (x). Hence, the Z-score represents the standard deviation that the sequence deviates from the mean MFE of the shuffled sequences. In the second equation, N refers to the number of randomly shuffled sequences, and R denotes the number of randomized sequences that have the MFE score lower than the original sequence (x).

##### 2.3.2 Ensemble-based features

Small RNAs vary in structures in vivo, and the distribution of these ensemble structures is modeled by the Boltzmann distribution. The calculation of various ensemble-based structure features was performed as described by Chan et al (Chan and Ding, 2008), that portray the global folding ability and cluster quality among the sRNA ensemble structures.

(1) Shannon base-pairing entropy measure (Shannon entropy). The shannon entropy is calculated from the ensemble of predicted optimal secondary structure. The equation is described as below.

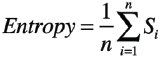

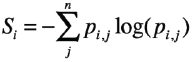

Where *P*_*i,j*_ the probability of base pair between nucleotides at sequence positions i and j, and n is the length of the sRNA sequence.

(2) Ensemble diversity (ensemble_diversity). The ensemble diversity means the base pair distance between the MFE and the other sub-optimal structures in the ensemble and obtained from “RNAfold -p” option.

(3) Frequency of the MFE structure in the ensemble (freq_mfe_ensemble). This value is also implemented in the RNAfold program, which indicates the uniqueness of the MFE structure in RNA ensemble.

(4) The free energy of the thermodynamic ensemble (free_energy_ensemble). This value is defined as the average free energy of all the suboptimal structure in the ensemble.

(5) Number of clusters (nclusters). This value means the numbers of clusters in the sRNA ensemble of structures found by RNACluster (Liu et al., 2008).

(6) Compactness. The compactness of each cluster was measured and further normalized by sequence length.

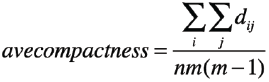

Where n is the number of the cluster, m is the number of structures within a cluster and *d*_*ij*_ is the base-pair distance.

(7) The number of high-frequency base-pairs in the ensemble (num_hifreq_ensemble). The base-pair frequency larger than 50% are defined as high-frequency base pairs.

(8) The average base-pair distance between the MFE structure and the suboptimal structure (ave_num_hifreq_percluster). Base-pair distance between two structures refers to the number of base pairs present in only one structure.

(9) Between-cluster sum of squares (BSS). The BSS value is a structural feature to describe the distance among the clusters of ensemble structures. The equation of calculation of BSS value is (4)

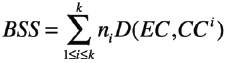

Where K is the optimal number of clusters, *n*_*i*_, is the number of structures in the ith cluster, D means the base-pair distance between two structures, is the ensemble centroid, and is the centroid of the ith cluster.

(10) Within cluster sum of squares (WSS). The WSS value is a structural measurement to describe the compactness of clusters of structures.

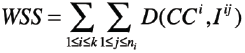

Where *I*^*ij*^ is the jth structure of the ith cluster, the other symbol is the same as the above equation.

##### 2.3.3 Structural statistics

Eighteen structural properties described by Tran et al. (Tran et al., 2009) have been used. They are divided into five types of structural elements measurement including stems, hairpin-loops, internal loops, bulges and multi-loops. The average length in each structural element along with the total counts and nucleotides of this structural element in MFE structure was measured.

#### 2.4 Genome sequences

I chose the genome sequences of four model organisms for genome-wide sRNA prediction. Two gram-negative strains were *Escherichia coli* MG1655 and *Salmonella* Typhimurium LT2, and two gram-positive strains were *Listeria monocytogenes* EGD-e and *Streptococcus agalactiae* NEM316. The genome sequences and annotations were downloaded from NCBI Genbank ftp site (http://www.ncbi.nlm.nih.gov/genbank/ftp/).

#### 2.5 Sliding-window approaches

I have used a sliding-window approach for genome-wide prediction of sRNAs. The initial window size was set to 160bp, which is the average small RNA size in BSRD database, with step size of 50bp. In each sliding-window, previously identified significant 17 sub structure features were detected, plus 20 sequence statistics were calculated.

#### 2.6 Machine learning implementation

The features ranking was performed using information gain evaluator with default parameters and 10-fold cross validation. The random forest, the C4.5 decision tree and neural network model were trained using WEKA (Frank et al., 2004). These models were combined with Adaboost for meta analysis with 10-fold cross validation. The neural network was trained using multiplayer Perceptron model (options: -L 0.3 -M 0.2 -N 500 -V 0 -S 0 -E 20 -H a). The other two classifiers considered were (i) the random forest classifier, trained with Adaboost (options: AdaBoostM1 -I 50 -K 0 -S 1) and (ii) the C4.5 decision tree classifier with default parameters.

#### 2.7 Performance analysis

The prediction accuracy was evaluated using various criteria including sensitivity (SN), specificity (SP), and F-measure. The model’s performance was evaluated using 10-fold crossvalidation. The receiver operating characteristic (ROC) curve was obtained by plotting true positive rate (SN) against the false positive rate (1-SP). The area under the ROC curve (AUC) was also calculated. They are defined as below:

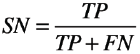

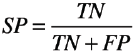

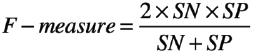

where TP, TN, FP, and FN denotes the numbers of true positives, true negatives, false positives, and false negatives, respectively.

#### 2.8 Statistical analyses

Numerous features of different groups were statistically analyzed using the R program (version 2.15.2). Values in different groups were compared using the paired or unpaired t test, and ANOVA with F test.

